# Distinctive retinal peri-arteriolar versus peri-venular amyloid plaque distribution correlates with the cognitive performance

**DOI:** 10.1101/2024.02.27.580733

**Authors:** Oana M. Dumitrascu, Jonah Doustar, Dieu-Trang Fuchs, Yosef Koronyo, Dale S. Sherman, Michelle Shizu Miller, Kenneth O. Johnson, Roxana O. Carare, Steven R. Verdooner, Patrick D. Lyden, Julie A. Schneider, Keith L. Black, Maya Koronyo-Hamaoui

## Abstract

**Introduction:** The vascular contribution to Alzheimer’s disease (AD) is tightly connected to cognitive performance across the AD continuum. We topographically describe retinal perivascular amyloid plaque (AP) burden in subjects with normal or impaired cognition.

**Methods:** Using scanning laser ophthalmoscopy, we quantified retinal peri-arteriolar and peri-venular curcumin-positive APs in the first, secondary and tertiary branches in twenty-eight subjects. Perivascular AP burden among cognitive states was correlated with neuroimaging and cognitive measures.

**Results:** Peri-arteriolar exceeded peri-venular AP count (p<0.0001). Secondary branch AP count was significantly higher in cognitively impaired (p<0.01). Secondary small and tertiary peri-venular AP count strongly correlated with clinical dementia rating, hippocampal volumes, and white matter hyperintensity count.

**Discussion:** Our topographic analysis indicates greater retinal amyloid accumulation in the retinal peri-arteriolar regions overall, and distal peri-venular regions in cognitively impaired individuals. Larger longitudinal studies are warranted to understand the temporal-spatial relationship between vascular dysfunction and perivascular amyloid deposition in AD.

**Highlights:**

- Retinal peri-arteriolar region exhibits more amyloid compared with peri-venular regions.
- Secondary retinal vascular branches have significantly higher perivascular amyloid burden in subjects with impaired cognition, consistent across sexes.
- Cognitively impaired individuals have significantly greater retinal peri-venular amyloid deposits in the distal small branches, that correlate with CDR and hippocampal volumes.

## Introduction

The vascular contribution to Alzheimer’s disease (AD) and AD-related dementias is increasingly recognized (1). The role of various vascular derangements in neurodegeneration had been underscored by multiple studies (2–10) and disruption of the blood-brain barrier proposed to be an early biomarker of cognitive dysfunction in AD (1). As retina is a central nervous system organ amenable to non-invasive and reproducible high-resolution imaging, whose blood-retina barrier shares similar pathophysiology to the blood-brain barrier (11–14), multiple static and dynamic retinal vascular biomarkers were investigated across the AD spectrum (11, 15–19). Retinal vascular pathology in AD was characterized using various retinal imaging technologies in humans, such as color and autofluorescence fundus photography, optical coherence tomography angiography (OCTA) (20–22), fluorescein angiography (23, 24), and targeted retinal pericyte imaging in animal models (25). On fundus photography, several vascular abnormalities had been reported in AD, such as venular narrowing, diminished vascular branching, increased tortuosity, and decreased arterial fractal dimension (19, 26–29) leading to the recent development of retinal photography-based deep learning algorithms to cost-effectively screen for AD in community settings (30, 31). Furthermore, retinal vascular tortuosity was proposed to improve the detection of the cerebral amyloid status as determined by 8F-florbetaben PET (32). Amyloid β-protein (Aβ) is an early core biomarker of AD, a prerequisite for AD diagnosis, and the target of AD-specific therapies (33), including the recently approved anti-Aβ monoclonal antibodies that bind with high affinity to Aβ plaque and/or soluble protofibrils (34, 35). These immune-based therapies showed efficacy in patients with mild cognitive impairment (MCI) and mild AD dementia (34–37).

Accurate detection of vascular-associated amyloid status in early AD remains an urgent unmet need in current clinical practice. The cost-effective and non-invasive detection of retinal amyloid burden carries the potential for early AD identification and monitoring (2, 38–40). Thus, several retinal amyloid imaging methodologies have recently been developed (41, 42), such as scanning laser ophthalmoscopy (SLO) (17, 39, 42–46) and hyperspectral retinal imaging (13, 39–41). SLO imaging offers the opportunity to visualize and quantify not only the Aβ plaques (AP), but also the retinal arteries and veins (17, 44, 45, 47). We have previously shown that retinal vascular geometric features correlate with retinal amyloid deposits, retinal venular tortuosity could discriminate between patients with normal and impaired cognition, and the discrimination is stronger when an index combining the AP count in the retinal proximal mid-periphery and retinal venular tortuosity is utilized (17). Moreover, our group has identified the significant accumulation of vascular and perivascular Aβ deposition in postmortem retinas of MCI (due to AD) and AD patients (16, 42, 48). We further showed that increased histopathological arteriolar Aβ_40_ deposition and tight junction loss in the inner blood-retinal barrier of prodromal and symptomatic AD patients are tightly correlated with cerebral amyloid angiopathy (CAA) severity and other AD related brain neuropathological changes (49).

Although various vascular alterations were described in retinal imaging studies in AD, it is currently unclear what type of vascular dysfunction, arteriolar or venular, predominates in preclinical or minimally symptomatic AD stages. It is also uncertain if retinal arteriolar or venular clearance limitations lead to AP deposition in the vascular-adjacent zones (termed here as perivascular AP), and how these evolve with AD progression. To elucidate the interactions between retinal vascular dysfunction and retinal AP deposition in early AD, we designed an exploratory analysis of a cohort of subjects with normal or impaired cognition that underwent retinal curcumin fluorescence amyloid imaging with SLO. We aimed to study the topographic relationship between perivascular AP deposition and retinal vessel types, to identify if retinal AP distribution is predominantly peri-arteriolar or peri-venular, in the proximity to the first-order vessels or further distal after the secondary or tertiary vessel bifurcation. We hypothesized that increased perivascular retinal AP count correlates with cognitive decline, and like in the AD brain, retinal AP deposits are more numerous in the peri-arteriolar than the peri-venular space. We compared the perivascular AP distribution in subjects with and without cognitive impairment and assessed the correlation between retinal peri-arteriolar and peri-venular AP count with cognitive and neuroimaging measures.

## Methods

### Study design

The prospective cohort study was approved by the Cedars-Sinai Medical Center Institutional Review Board (IRB) and was performed in accordance with the ethical standards as laid down in the 1964 Declaration of Helsinki and its later amendments or comparable ethical standards. We enrolled subjects older than 40 years of age with subjective cognitive decline that provided written informed consent prior to the enrollment. Exclusion criteria were prespecified, limited to a history of glaucoma, allergy to mydriatic eye drops, curcumin, or vitamin E. All subjects underwent retinal fluorescence imaging with a confocal scanning laser ophthalmoscope (SLO Retia^TM^, CenterVue SpA; Figure 1A) that utilizes blue light for the excitation of curcumin emission to obtain fluorescent images of the retina (Figure 1B-C), following a previously reported study design (17, 42, 44). They also underwent standard neuropsychological testing conducted by a licensed neuropsychologist (DS), that included the Montreal Cognitive Assessment (MOCA), global Clinical Dementia Rating (CDR), general cognitive (ACS-test of Premorbid Functioning) and specific cognitive domain assessments: attention and concentration (Wechsler Adult Intelligence Scale (WAIS)-IV); verbal memory (California Verbal Learning Test (CVLT) II, Wechsler Memory Scale (WMS)-IV, and Logical Memory II); non-verbal memory (Rey Complex Figure Test and Recall (RCFT) 30 min, and Brief Visuo-Spatial Memory Test Revised (BVMT-R) Delayed Recall); language (Fluency-Letter (FAS) and Fluency-category (animals)); visuo-spatial ability (Rey Complex Figure Test and Recognition Trial (RCFT) Copy); speed of information processing (Trails A and B); and symptom validity and functional status (SF-36 Physical Component Score (PCS) and Mental Component Score (MCS)). The subject’s emotional status was assessed using the Beck Depression Inventory II, Geriatric Depression Scale, and Profile of Mood State/Total Mood Disturbance. All subjects underwent 3 Tesla non-contrast structural brain magnetic resonance imaging (MRI). Automated NeuroQuant software was used for brain volumetric analysis (50) and the following parameters were collected: total intracranial volume (ICV) (cm^3^), hippocampal volume (HV) (cm^3^), and inferior lateral ventricle volume (ILVV) (cm^3^). The number and volume of white matter hyperintensities (WMHI) were measured using SPIN Software (SpinTech, Bingham Farms, MI).

**Figure 1.**
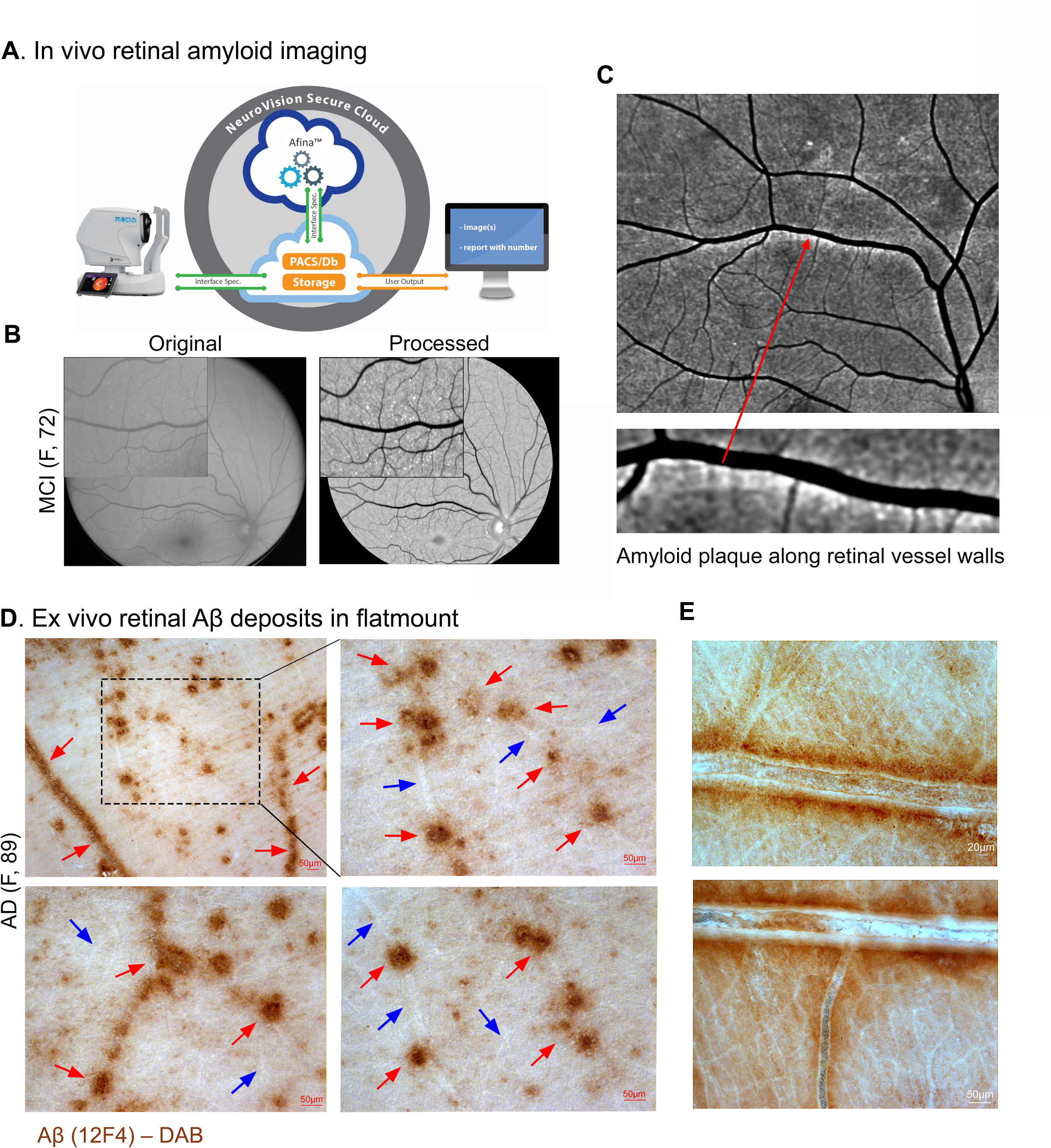
Retinal vascular amyloid imaging. (**A**) Pipeline of fluorescent imaging using Retia® SLO, Afina^TM^ cloud storage, and a fully automated image processing and analyses. (**B**) Representative retinal fluorescent image of before and after image processing. (**C**) Representative image of an AD patient demonstrating retinal amyloid deposits (white) along the blood vessels. (**D**) Representative images from a confirmed AD patient. Postmortem retinal flatmount immunolabeled with anti-c monoclonal antibody (12F4; brown) and peroxidase-base 3,3’ diaminobenzidine (DAB) immunostaining. Typical Aβ plaque structures and Aβ deposition along and inside blood vessels (red arrows) and ‘plaque-free’ regions of retinal blood vessels (blue arrows). (**E**) High magnification images of perivascular and vascular Aβ deposits in retinal flatmounts of an AD patient.

### Aβ immunostaining in human retinal flatmounts

Eyes from a deceased AD female patient was obtain from Rush University’s Alzheimer’s disease research center (ADRC ORA# 18011111) and processed for retinal flatmounts and stained for Aβ_42_ as previously described (13, 42, 51). Briefly, the neurosensory retina was isolated from the whole eye and the vitreous thoroughly removed before flatmount preparation. The retina was then treated with antigen retrieval (pH 6.1; S1699, DAKO) followed by peroxidase-base 3,3’ diaminobenzidine (DAB) immunostaining with anti-Aβ_42_ antibodies (12F4; Biolegend # 805501).

Bright-field images were acquired using a Carl Zeiss Axio Imager Z1 fluorescence microscope (Carl Zeiss MicroImaging, Inc.) equipped with ApoTome, AxioCam MRm and AxioCam HRc cameras (Figure 1D-E; Supplementary Figure 1). Histological studies were performed under the IRB protocol Pro00055802 at Cedars-Sinai Medical Center.

### Human retinal imaging analysis

Retinal AP imaging analysis was performed as per our previously reported methodology in a population of mostly Caucasian subjects (17, 42, 44), that was recently replicated in a Japanese population (45). The researchers conducting the retinal images processing and AP quantification were blinded to the patients’ clinical characteristics. Retinal fluorescence images from the left eye supero-temporal quadrant were used for the perivascular AP identification and quantification. For this purpose, the retinal arteries and veins were manually identified by a trained observer based on their vessel diameter (vein > artery (52)), location, and morphology, and traced using Adobe Photoshop CC 2020. Vessels stemming from the optic disc were defined as the “primary (1°) main branch” for each venular and arteriolar network. After the defined main primary branch bifurcation, the subsequent divisions were called the “secondary (2°) main branches” for all venules and arterioles. After these main secondary arterioles and venules split, the following divisions were called “tertiary (3°) branches” (Figure 2C-D). All protruding vessels from main branches were deemed ‘small’. For example, a protruding vessel from the main primary branch was called “primary branch-small”. If they were protruding from the main secondary branch, they were considered “secondary branch - small” (Figure 2C’-D’). Tertiary vessels and their branches had relatively equal diameters, therefore all vessels following the second bifurcation were grouped as tertiary branches. For analytical purposes, main branches along with protruding vessels were additionally compiled into one measurement, deemed primary branches and secondary branches, encompassing both main and small primary and respectively secondary branches.

**Figure 2.**
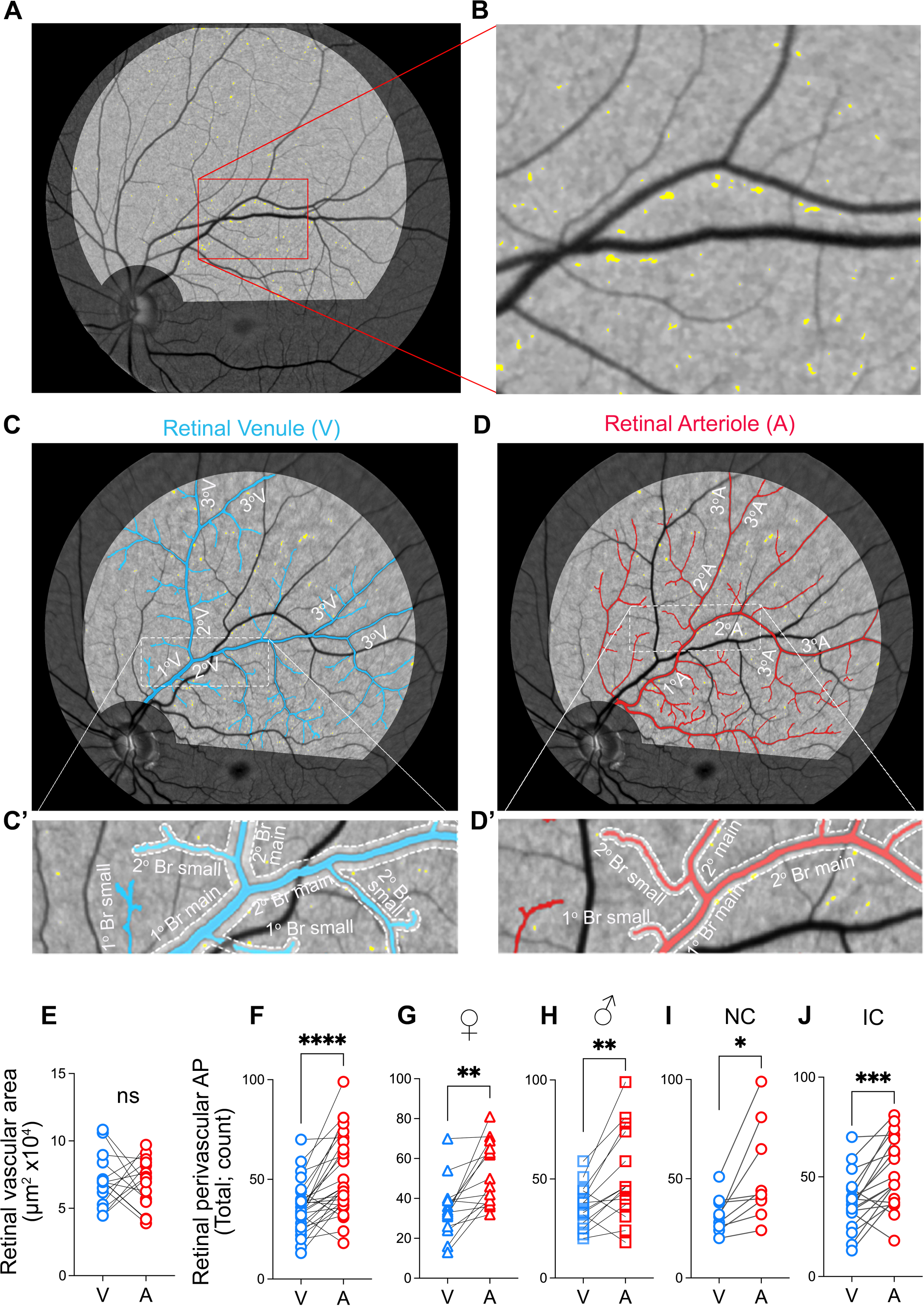
Retinal peri-venular and peri-arteriolar amyloid plaque distribution. **(A-B)** Representative retinal fluorescent fundus image illustrating retinal curcumin-positive amyloid hyperfluorescent plaques in the left eye supero-temporal quadrant (**A**, magnification **B**). Illustration of the primary, secondary and tertiary retinal venular (**C**) and arteriolar (**D**) branches. Magnifications of the peri-venular (**C’**) and peri-arteriolar (**D’**) area used for the amyloid plaque (AP) quantification; boundary zone delineated by dotted lines producing a perivascular area of one equivalent vessel diameter on either side. **(E)** Quantitative analysis of retinal perivascular area stratified by venules (V) and arterioles (A) showing no significant difference between the total area for each vessel type in the analyzed supero-temporal region. (**F-J)** Quantitative analyses of retinal perivascular AP count stratified by V versus A in the total branches (**F**), in females (**G**), males (**H**), individuals with normal cognition (**I**) and impaired cognition (**J**). Individual data points are shown. * P<0.05, ** P<0.01, *** P< 0.001, **** P< 0.0001 by paired two-tailed Student’s t test. NC, Normal cognition; IC, Impaired cognition; 1°V, primary venular branch; 2°V, secondary venular branch; 3°V tertiary venular branch; 1°A, primary arteriolar branch, 2°A, secondary arteriolar branch; 3°A, tertiary arteriolar branches. Color code vessel type: blue – peri-venular; red – peri-arteriolar.

Starting within the highlighted imaging field and closest to the optic disc, we measured the diameter of the primary vessel at pre-set intervals along its length using the line segment tool. We calculated the diameter of each vessel by averaging these measurements from pre-set intervals as well as most proximal and distal segments. We next manually traced through the middle of the vessel using the curvature tool, emphasizing accuracy. We set the diameter of the drawn external perivascular line to automatically display three times the diameter of the vessel, producing a “perivascular area” of one equivalent vessel diameter on either side. We edited the line opacity to 40% so that plaque positive signals can be detected within this perivascular area. The plaques that touched the border were counted as perivascular, even if their entire body was not within the boundary. We repeated these steps for all venules and arterioles to trace all peri-arteriolar and peri-venular areas. We counted all hyperfluorescent AP positive signals that fall within the boundaries of the perivascular areas and separated them by 1) vessel type (peri-venular versus peri-arteriolar), and 2) location (primary vessel, primary branch, secondary vessel, secondary branch, or tertiary vessel).

### Statistical analysis

Descriptive statistics were performed, and continuous demographic and clinical data are presented as the mean ± standard deviation (SD) in the text and tables. Analyses comparing peri-arteriolar vs peri-venular AP count utilized paired and unpaired, two-tailed Student’s t tests or one-way analysis of variance (ANOVA) followed by Tukey’s multiples comparisons test. Subjects were partitioned into three groups according to the Clinical Dementia Rating (CDR) (0.5, questionable impairment; 1, mild cognitive impairment; and 2, moderate cognitive impairment) (53). They were also dichotomized using MOCA cut-off of 26 (54) and the neuropsychometric diagnosis (normal cognition versus impaired cognitive performance). Group differences between continuous variables were conducted using Student’s t-test, while nominal qualitative variables were assessed using chi-square test. Differences in continuous variables between levels of CDR were assessed through ANOVA with Tukey’s test to correct for multiple comparisons. Pearson’*s* r correlation analysis was conducted to assess the relationship between retinal perivascular AP count and cognitive and brain imaging volumetric measures. All statistical analyses were performed using GraphPad Prism 8.3 with two-sided tests and a significance level of < 0.05.

## Results

A total of 34 patients underwent retinal and brain imaging, and neuropsychometric evaluation. Noninvasive retinal amyloid images were acquired in predefined geometrical regions using the Retia^TM^ SLO, followed by a fully automated Afina^TM^-based cloud storage and Neurovision imaging-based image processing output (Figure 1A). Representative retinal curcumin fluorescence imaging detected multiple retinal hyperfluorescence spots (APs) that were more visible after image processing (Figure 1B, white dots). APs were also detected along retinal blood vessels (Figure 1C). Ex vivo immunohistochemical analysis of Aβ in postmortem retinal flatmounts from an 89-years-old female with AD is presented to illustrate multiple abluminal and perivascular AP, accumulating along and inside retinal blood vessels (Figure 1D-E).

Perivascular AP analyses included twenty-eight patients that had the left eye supero-temporal retinal images suitable for the vascular tracing and perivascular AP quantitative analysis. Mean (SD) age was 65 (7.4) years, and 50% were female. Mean (SD) MOCA was 25 (5.6) and median MOCA was 27 (range 4–32). Eleven subjects had a CDR of 0.5, 14 had a CDR of 1, and 3 had a CDR of 2. Based on formal neuropsychometric cognitive evaluation, 9 (32.1%) patients had normal cognition (NC) and 19 (67.9%) had impaired cognition (IC; six amnestic MCI, nine multidomain MCI, one non-amnestic MCI, and three probable AD). The cohorts with normal and impaired cognition were matched for age, sex, and years of education (Table 1).

**Table 1.** Cohort Demographic and Cognitive Characteristics.

| **Demographics** | **All (n=28)** | **Normal Cognition (n=9)** | **Impaired Cognition (n=19)** | **P-value** |
| --- | --- | --- | --- | --- |
| Average age (SD) | 65 (7.4) | 65 (5.3) | 65 (8.3) | 0.83 |
| Sex (male n (%)) | 14 (50%) | 4 (44.4%) | 10 (52.6%) | – |
| Years of Education Mean (SD) | 17 (1.8) | 17 (1.5) | 17 (1.9) | 0.94 |
| **Neurocognitive characterization** | **All (n=28)** | **MOCA >26 (n=15)** | **MOCA ≤26 (n=13)** |  |
| Average age (SD) | 65 (7.4) | 64 (7.2) | 66 (7.6) | 0.36 |
| Sex (male n (%)) | 14 (50%) | 6 (40%) | 8 (60%) | – |
| Years of Education Mean (SD) | 17 (1.8) | 17 (1.5) | 17 (2.1) | 0.69 |
|  | **Score** | **Total** | **Female** | **Male** |
| **Standard Global CDR (n =28)** | 0.5 | 11 | 7 | 4 (36.4%) |
|  | 1 | 14 | 6 | 8 (57.1%) |
|  | 2 | 3 | 1 | 2 (66.7%) |

NC – normal cognition, IC – impaired cognition (refers to patients with diagnosis of mild cognitive impairment
(aMCI) and Alzheimer’s disease (AD)), MOCA – Montreal cognitive assessment, CDR – Clinical dementia rating.
Statistical analysis was established using Student’s t test.

Despite no significant difference between the perivascular area of retinal venules and arterioles (Figure 2E), the total retinal perivascular AP count was significantly greater in the peri-arteriolar (A) compared to peri-venular (V) area (Figure 2F; A: 51.96 ± 19.65 versus V: 35.57 ± 12.66, P<0.0001; extended data for each vascular subtype in Supplementary Table 1). These results remained consistent irrespective of the sex (Figure 2G-H, Supplementary Table 2) and the degree of cognitive impairment (Figure 2I-J). Notably, the increased retinal peri-arteriolar versus peri-venular AP count is statistically more significant in individuals with impaired cognition (IC) compared with those with normal cognition (NC) (Fig 2I versus 2J). In contrary, only in the retinal primary main branches, the peri-venular AP count was greater than the peri-arteriolar plaque count (P=0.013; Supplementary Table 1). After the first bifurcation, retinal peri-arteriolar AP burden remained significantly greater than the peri-venular AP burden at all levels, aside for a similar non-significant trend in the tertiary branches (Supplementary Figure 2A-E, Supplementary Table 1). When stratified by cognitive status, IC compared to NC subjects had significantly greater retinal perivascular AP burden in the secondary branches (P=0.0037), both in the total peri-venular (P= 0.0011) and total peri-arteriolar areas (P=0.015), which was substantially (2.4-2.9-fold) greater for the secondary small branches (Figure 3A-E and Table 2; extended data in Supplementary Figure 2F-H). Tertiary branches also exhibited significantly greater retinal AP burden in the peri-venular areas in the IC group (P=0.031, Supplementary Figure 2H). Interestingly, the main primary branches showed non-significant trend of greater perivascular AP burden in NC versus IC subjects (P=0.08, Table 2).

**Figure 3.**
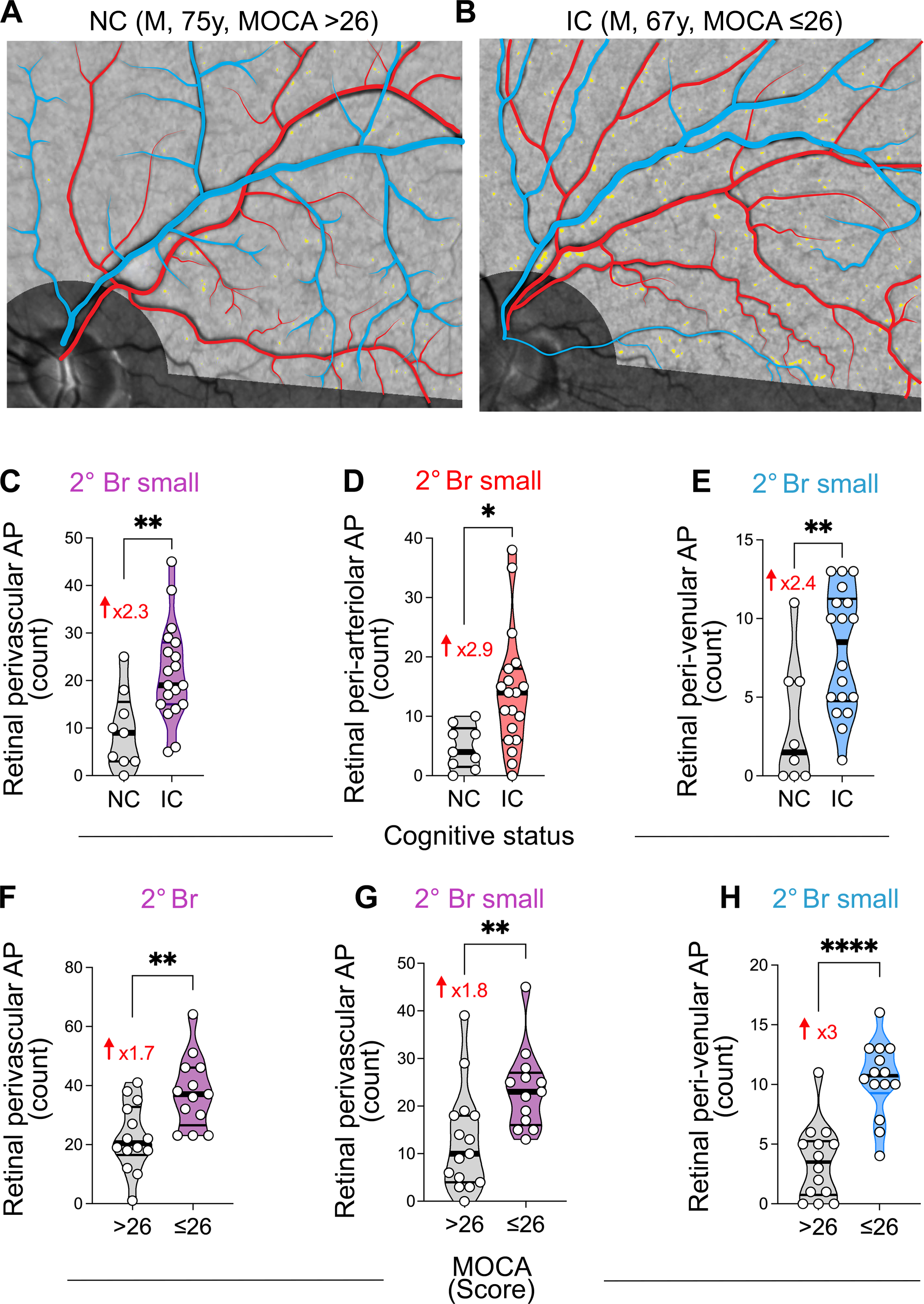
Retinal peri-arteriolar and peri-venular amyloid plaque count stratified by cognitive status and MOCA scores. (**A-B**) Representative fundus images of retinal perivascular amyloid plaque (AP) showing their density and distribution along arterioles (red tracing) and venules (blue tracing), in individuals with normal cognition (NC; **A**) or impaired cognition (IC; **B**). (**C-E**) Quantitative analyses of retinal AP count stratified by cognitive status, NC versus IC, in perivascular (**C**), peri-arteriolar (**D**), and peri-venular (**E**), for the secondary (2°) small branches. (**F-H**) Quantitative analyses of AP count stratified by MOCA scores of 26 or lower compared with greater than 26, in perivascular 2° branches (**F**), 2° small branches (**G**), and peri-venular 2° small branches (**H**). Violin plots are showing individual data points, median and interquartile range. Statistics: * P<0.05, ** P<0.01, **** P< 0.0001, by unpaired two-tailed Student’s t test. M, male; MOCA, Montreal Cognitive Assessment; y, years. Color code for vessel type: purple – perivascular; blue – peri-venular; red – peri-arteriolar.

**Table 2.** Retinal perivascular amyloid plaques (AP) in cognitively normal and impaired subjects.

| DIAGNOSTIC GROUP |  | NC |  |  | IC |  |  | Fold Change | Unpaired t test |
| --- | --- | --- | --- | --- | --- | --- | --- | --- | --- |
| Vascular type |  | Mean | SD | n | Mean | SD | n | FC | P value |
| Perivascular AP | **Total** |  |  | 9 |  |  | 19 |  |  |
|  | **Primary Br (total)** | 43.56 | 35.12 | 9 | 24.32 | 16.85 | 19 | 0.56 | 0.058 |
|  | *Primary Br – main* | 12.33 | 7.28 | 9 | 7.32 | 6.60 | 19 | 0.59 | 0.080 |
|  | *Primary Br – small* | 31.22 | 29.55 | 9 | 17.00 | 11.64 | 19 | 0.54 | 0.076 |
|  | **Secondary Br (total)** | 14.67 | 10.71 | 9 | 19.78 | 10.47 | 19 | 1.81 | **0.0037** |
|  | *Secondary Br – main* | 10.33 | 5.89 | 9 | 14.26 | 6.36 | 19 | 1.38 | 0.13 |
|  | *Secondary Br – small* | 9.44 | 8.14 | 9 | 21.53 | 10.15 | 19 | 2.28 | **0.0044** |
|  | **Tertiary Br** | 19.56 | 12.63 | 9 | 27.89 | 19.33 | 19 | 1.43 | 0.25 |
| Peri-Venular AP | **Total** |  |  | 9 |  |  | 19 |  |  |
|  | **Primary Br (total)** | 18.67 | 13.17 | 9 | 11.68 | 9.15 | 19 | 0.63 | 0.11 |
|  | *Primary Br – main* | 7.89 | 4.76 | 9 | 4.63 | 5.05 | 19 | 0.59 | 0.12 |
|  | *Primary Br – small* | 10.78 | 9.67 | 9 | 7.05 | 5.65 | 19 | 0.65 | 0.21 |
|  | **Secondary Br (total)** | 6.25 | 4.59 | 8 | 13.39 | 4.53 | 18 | 2.14 | **0.0011** |
|  | *Secondary Br – main* | 3.11 | 1.90 | 9 | 5.44 | 2.999 | 18 | 1.75 | **0.044** |
|  | *Secondary Br – small* | 3.25 | 4.03 | 8 | 7.94 | 3.90 | 18 | 2.44 | **0.0098** |
|  | **Tertiary Br** | 4.43 | 2.76 | 7 | 11.93 | 8.23 | 15 | 2.69 | **0.031** |
| Peri-Arteriolar AP | **Total** |  |  | 9 |  |  | 19 |  |  |
|  | **Primary Br (total)** | 24.89 | 24.43 | 9 | 12.63 | 11.96 | 19 | 0.51 | 0.083 |
|  | *Primary Br – main* | 4.44 | 4.13 | 9 | 2.68 | 2.77 | 19 | 0.60 | 0.19 |
|  | *Primary Br – small* | 23.00 | 21.27 | 8 | 10.50 | 10.14 | 18 | 0.46 | 0.051 |
|  | **Secondary Br (total)** | 12.00 | 5.70 | 9 | 23.11 | 12.13 | 19 | 1.93 | **0.015** |
|  | *Secondary Br – main* | 7.22 | 4.35 | 9 | 9.11 | 4.43 | 19 | 1.26 | 0.30 |
|  | *Secondary Br – small* | 4.78 | 3.60 | 9 | 14.00 | 10.01 | 19 | 2.93 | **0.013** |
|  | **Tertiary Br** | 14.44 | 9.89 | 9 | 16.56 | 10.94 | 18 | 1.15 | 0.63 |

AP – Amyloid plaque; Br – Branch; FC – Fold change; IC – Impaired cognition; NC – Normal cognition; SD – Standard deviation. Statistical analysis was established using Student's t test. \*P < 0.05, \*\*P < 0.01.

When compared to subjects with MOCA greater than 26 (Figure 3F-H and Table 3; extended data in Supplementary Figure 2I), subjects with MOCA scores 26 or lower had significantly greater total secondary branch perivascular AP count (P=0.002), with the most substantial 3-fold increase in the peri-venular small branch areas (3.50 ± 3.13 vs 10.46 ± 3.26, P<0.0001).

**Table 3.**
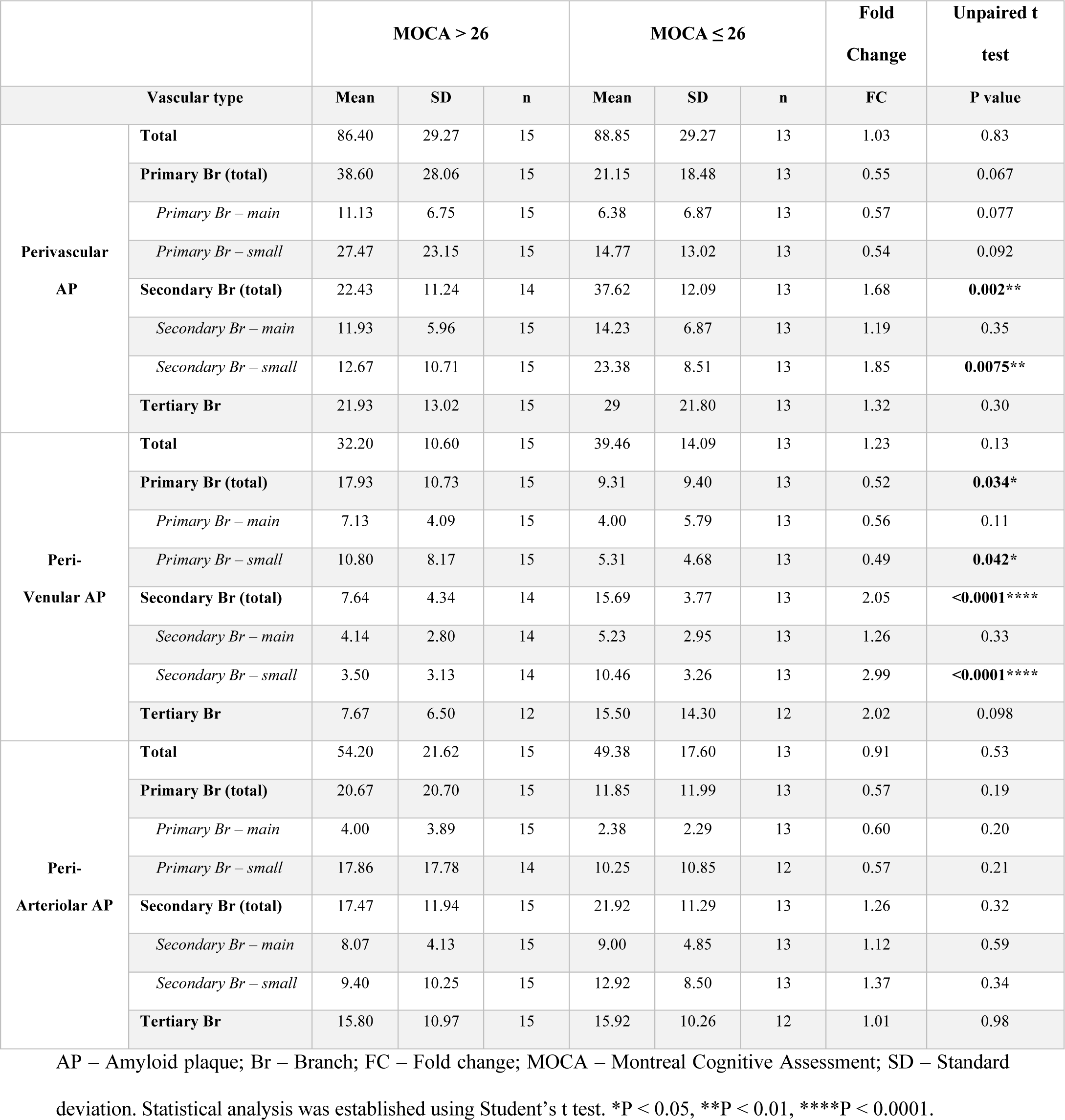
Retinal perivascular amyloid plaques (AP) separated by MOCA score.

Patients with high CDR scores (Supplementary Figure 3 and Supplementary Table 3) exhibited greater total retinal perivascular AP count in the secondary branches (P=0.020), and significantly higher values in tertiary branch areas on further topographic analysis (P=0.0058; Supplementary Figure 3A-B). Total peri-venular AP burden, and especially in the tertiary branches, was significantly greater in subjects with higher CDR (P=0.0025 and P=0.0007 respectively; Supplemental Figure 3C-D). Peri-arteriolar secondary branch total regions, and both main and small branches, also showed significantly greater AP count in subjects with CDR 2 compared to CDRs 0.5 and 1 (Supplementary Figure 3E and Supplementary Table 3).

Pearson’s correlation analyses indicated that retinal perivascular AP count correlated with the CDR scores, with the most significant correlation in this cohort observed for the total peri-venular AP count (Figure 4A and Table 4; *r*=0.66, P=0.0001). Topographically, the tertiary branch peri-venular AP count (*r*=0.61, P=0.0016) and the secondary branch peri-arteriolar AP count (*r*=0.51, P=0.0055) showed significant correlation with CDR (Table 4). The secondary small branch peri-venular AP count showed the highest correlation with the MOCA score (Figure 4B; *r*=-0.51, P=0.0063).

**Figure 4.**
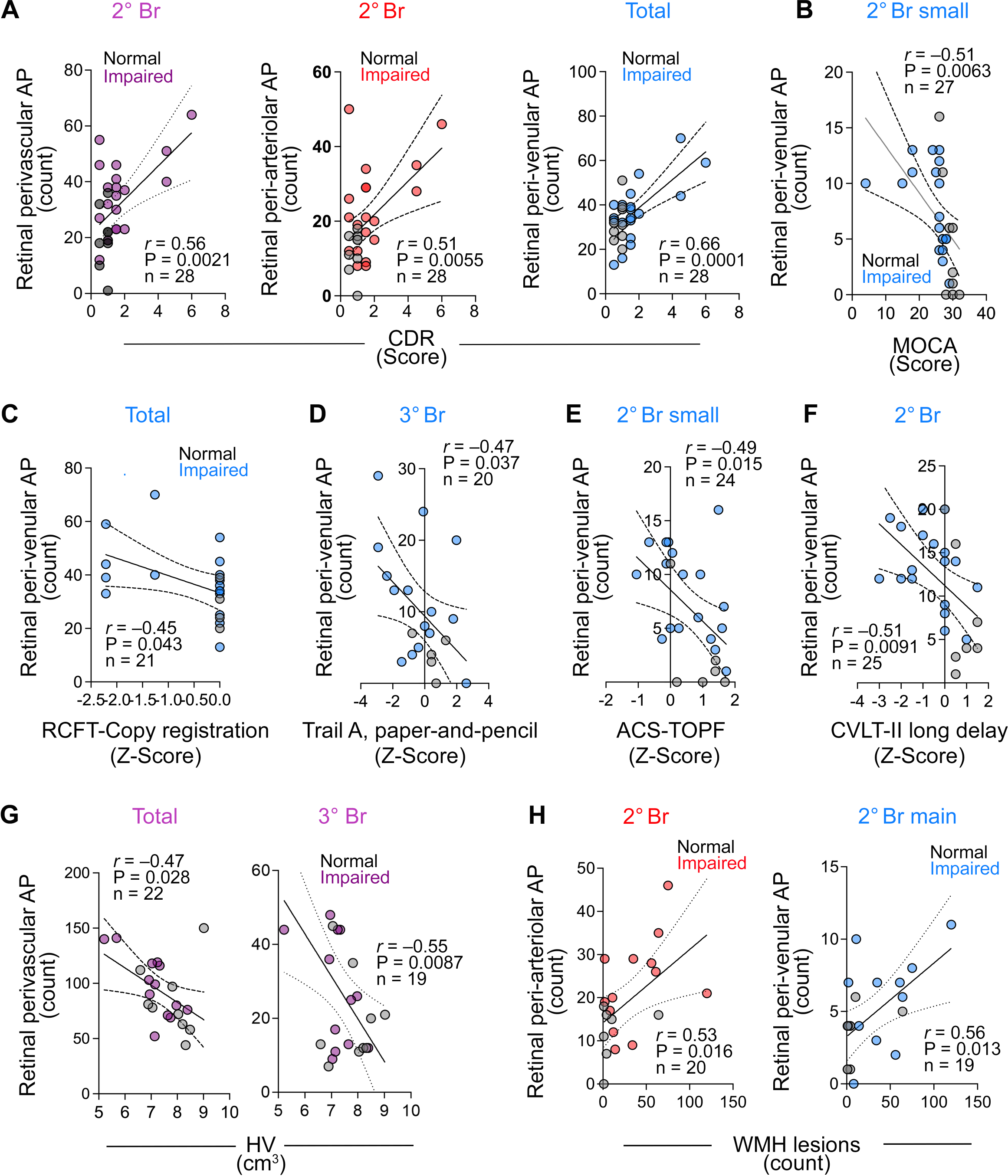
Correlations between retinal perivascular amyloid plaque distribution with cognitive and neuroimaging measures. Pearson’s *r* correlation analyses between retinal perivascular AP count and CDR (**A**), MOCA (**B**), RCFT-copy registration (**C**), Trail A-paper and pencil (**D**), ACS-TOFF (**E**), CVLT-II Long delay (**F**), hippocampal volume (**G**) and white matter hyperintensities lesions (**H**). AP, Amyloid plaques; CDR, Clinical dementia rating; MOCA, Montreal cognitive assessment; RCFT, Rey Complex Figure Test and Recognition Trial; ACS-TOPF, test of Premorbid Functioning; CVLT, California Verbal Learning Test; HV, hippocampal volume, WMH, White matter hypertensity; 2° Br, secondary branches; 3° Br, tertiary branches. Color code for vessel type: purple – perivascular; blue – peri-venular; red – peri-arteriolar.

**Table 4.** Pearson’s *r* correlations between peri-venular or peri-arteriolar AP with cognitive status and brain imaging parameters.

|  |  | MOCA<br>n = 24-28 |  | CDR<br>n = 24-28 |  | HV<br>n = 19-22 |  | Number of WMHI<br>n = 17-20 |  |
| --- | --- | --- | --- | --- | --- | --- | --- | --- | --- |
| Vascular type | | $r$ | P | $r$ | P | $r$ | P | $r$ | P |
| Perivascular<br>AP | **Total** | 0.03 | ns | **0.46** | **0.01** | **-0.47** | **0.028** | 0.34 | ns |
|  | **Primary Br (total)** | 0.29 | ns | -0.15 | ns | 0.09 | ns | -0.24 | ns |
|  | *Primary Br – main* | 0.19 | ns | 0.04 | ns | -0.20 | ns | 0.01 | ns |
|  | *Primary Br – small* | 0.30 | ns | -0.21 | ns | 0.19 | ns | -0.31 | **0.045** |
|  | **Secondary Br (total)** | **-0.38** | **0.05** | **0.56** | **0.0021** | -0.41 | ns | **0.56** | **0.010** |
|  | *Secondary Br – main* | -0.11 | ns | **0.40** | **0.034** | -0.27 | ns | **0.48** | **0.033** |
|  | *Secondary Br – small* | **-0.42** | **0.02** | **0.49** | **0.0078** | -0.40 | ns | **0.45** | **0.047** |
|  | **Tertiary Br** | -0.07 | ns | **0.53** | **0.0040** | **-0.55** | **0.0087** | 0.44 | 0.052 |
| Peri-Venular AP | **Total** | -0.26 | ns | **0.66** | **0.0001** | **-0.51** | **0.016** | 0.40 | ns |
|  | **Primary Br (total)** | 0.27 | ns | -0.06 | ns | 0.02 | ns | -0.33 | ns |
|  | *Primary Br – main* | 0.20 | ns | 0.05 | ns | -0.15 | ns | -0.07 | ns |
|  | *Primary Br – small* | 0.27 | ns | -0.12 | ns | 0.14 | ns | **-0.45** | **0.045** |
|  | **Secondary Br (total)** | **-0.44** | **0.022** | 0.31 | ns | -0.24 | ns | 0.43 | ns |
|  | *Secondary Br – main* | -0.03 | ns | 0.21 | ns | -0.34 | ns | **0.56** | **0.013** |
|  | *Secondary Br – small* | **-0.51** | **0.0063** | 0.25 | ns | -0.11 | ns | 0.15 | ns |
|  | **Tertiary Br** | -0.38 | ns | **0.61** | **0.0016** | **-0.51** | **0.026** | **0.52** | **0.032** |
| Peri-Arteriolar AP | **Total** | 0.21 | ns | 0.24 | ns | -0.37 | ns | 0.23 | ns |
|  | **Primary Br (total)** | 0.25 | ns | -0.19 | ns | 0.12 | ns | -0.14 | ns |
|  | *Primary Br – main* | 0.10 | ns | 0.02 | ns | -0.21 | ns | 0.11 | ns |
|  | *Primary Br – small* | 0.25 | ns | -0.16 | ns | 0.06 | ns | -0.13 | ns |
|  | **Secondary Br (total)** | -0.24 | ns | **0.51** | **0.0055** | -0.42 | ns | **0.53** | **0.016** |
|  | *Secondary Br – main* | -0.13 | ns | **0.42** | **0.026** | -0.19 | ns | 0.31 | ns |
|  | *Secondary Br – small* | -0.23 | ns | **0.43** | **0.022** | **-0.42** | **0.049** | **0.49** | **0.028** |
|  | **Tertiary Br** | 0.18 | ns | 0.19 | ns | **-0.46** | **0.033** | 0.13 | ns |

AP – Amyloid plaque; Br – Branch; CDR – Clinical dementia rating; HV – Hippocampal volume; MOCA – Montreal Cognitive Assessment; WMHI – White matter hypertensities.

The evaluation of specific cognitive domains (Figure 4, Table 5; extended data in Supplementary Figure 4) revealed greater overall retinal peri-venular AP accumulation in subjects with lower visuo-spatial ability Rey Complex Figure Test and Recognition Trial (RCFT)-Copy registration (Figure 4C; *r*=-0.45, P=0.043) and speed of information processing (Trail A paper and pencil; Table 5; *r*= −0.42, P=0.039) Z-scores. Similar moderate correlations were found between lower speed of information processing (Trail A paper and pencil) Z-scores and retinal AP counts in peri-venular tertiary branches (Figure 4D; r=-0.47, p=0.037) and peri-arteriolar secondary branches (Table 5; r=-0.44, P=0.027). We also noted higher retinal peri-venular AP count in distal secondary branches in subjects with lower Z-scores on the general cognitive Test of Premorbid Functioning ACS (Figure 4E; r=-0.49, *r*=0.015) and the California Verbal Learning Test (CVLT) II Long Delay (Figure 4F; *r*=-0.51, P=0.0091). Interestingly, positive correlations were found between peri-arteriolar primary branch AP count and RCFT recall 20min and CVLT-II Z-scores (*r*=0.41, P=0.039 and *r*=0.53, P=0.0052 respectively; Supplementary Figure 4A-B). Symptom validity and functional status (SF-36PCS) also positively correlated with retinal AP count in primary branches of the venular area only (Supplementary Figure 4C).

**Table 5.** Pearson’s *r* correlations between peri-venular or peri-arteriolar AP and cognitive domains.

|  |  | RCFT-Copy<br>(Z Score)<br>n = 18-21 |  | ACS-TOPF<br>(Z Score)<br>n = 21-25 |  | CVLT-II Long Delay<br>(Z-Score)<br>n = 22-26 |  | Trails A paper-and-pencil (Z-score)<br>n = 21-25 |  |
| --- | --- | --- | --- | --- | --- | --- | --- | --- | --- |
| Vascular type | | $r$ | P | $r$ | P | $r$ | P | $r$ | P |
| Perivascular AP | Total | -0.18 | ns | 0.07 | ns | -0.03 | ns | -0.26 | ns |
|  | Primary Br | 0.25 | ns | 0.20 | ns | **0.46** | **0.017** | 0.18 | ns |
|  | Primary Br – main | 0.09 | ns | 0.35 | ns | **0.43** | **0.028** | 0.11 | ns |
|  | Primary Br – small | 0.31 | ns | 0.13 | ns | **0.44** | **0.024** | 0.20 | ns |
|  | Secondary Br | -0.38 | ns | -0.27 | ns | **-0.41** | **0.036** | **-0.40** | **0.047** |
|  | Secondary Br – main | -0.31 | ns | -0.06 | ns | -0.10 | ns | **-0.47** | **0.02** |
|  | Secondary Br – small | -0.32 | ns | -0.31 | ns | **-0.47** | **0.014** | -0.25 | ns |
|  | Tertiary Br | -0.28 | ns | 0.06 | ns | **-0.40** | **0.046** | -0.30 | ns |
| Peri-Venular AP | Total | **-0.45** | **0.043** | -0.14 | ns | -0.21 | ns | **-0.42** | **0.039** |
|  | Primary Br | 0.16 | ns | 0.19 | ns | 0.28 | ns | 0.16 | ns |
|  | Primary Br – main | 0.15 | ns | 0.29 | ns | 0.27 | ns | 0.18 | ns |
|  | Primary Br – small | 0.14 | ns | 0.09 | ns | 0.24 | ns | 0.11 | ns |
|  | Secondary Br | -0.34 | ns | -0.38 | ns | **-0.51** | **0.0091** | -0.31 | ns |
|  | Secondary Br – main | 0.02 | ns | 0.07 | ns | -0.20 | ns | -0.32 | ns |
|  | Secondary Br – small | -0.44 | ns | **-0.49** | **0.015** | **-0.49** | **0.014** | -0.17 | ns |
|  | Tertiary Br | -0.40 | ns | -0.17 | ns | **-0.45** | **0.036** | **-0.47** | **0.037** |
| Peri-Arteriolar AP | Total | 0.05 | ns | 0.19 | ns | 0.09 | ns | -0.10 | ns |
|  | Primary Br | 0.24 | ns | 0.17 | ns | **0.50** | **0.010** | 0.15 | ns |
|  | Primary Br – main | -0.02 | ns | 0.32 | ns | **0.53** | **0.005** | -0.02 | ns |
|  | Primary Br – small | 0.24 | ns | 0.12 | ns | **0.44** | **0.030** | 0.13 | ns |
|  | Secondary Br | -0.30 | ns | -0.12 | ns | -0.34 | ns | -0.31 | ns |
|  | Secondary Br – main | -0.43 | ns | -0.10 | ns | -0.09 | ns | **-0.44** | **0.027** |
|  | Secondary Br – small | -0.17 | ns | -0.10 | ns | -0.37 | ns | -0.18 | ns |
|  | Tertiary Br | 0.06 | ns | 0.17 | ns | -0.32 | ns | -0.05 | ns |

AP – Amyloid plaque; Br – Branch; RCFT – Rey Complex Figure Test and Recognition Trial; ACS-TOPF – test of Premorbid Functioning; CVLT – California Verbal Learning Test.

Correlation analyses with neuroimaging measurements showed inverse relationships between HV and retinal AP count in perivascular total area (Figure 4G, left; *r*=-0.47, P=0.028), and moreover, tertiary branch area (Figure 4G, right; *r*=-0.55, P=0.0087). Comparable inverse correlations are found between HV and AP count in peri-venular total and tertiary branches (*r*=-0.51, P=0.016 and *r*=-0.51, P=0.026, respectively), as well as AP count in peri-arteriolar secondary small and tertiary branches (*r*=-0.42, P=0.049 and *r*=-0.46, P=0.033, respectively; Table 4 and Supplementary Figure 4D). In contrast, the number of WMHI (Figure 4H, Table 4, and Supplementary Figure 4E) positively correlated with retinal AP count in the peri-arteriolar secondary branch (*r*=0.53, P=0.016) and peri-venular secondary main and tertiary branches (*r*=0.56, P=0.013 and *r*=0.52, P=0.032 respectively). None of the perivascular AP counts showed significant correlations with total intracranial volumes, but some were shown with the volume of WMHI and subcortical Fazekas scores (Supplementary Figure 4F-G).

## Discussion

We describe the topographic distribution of retinal peri-arteriolar and peri-venular amyloid deposits in a pilot noninvasive retinal imaging study in living human subjects with normal or impaired cognition (mostly MCI). To our knowledge, this description has not been conducted before. Our exploratory analysis of the interaction between two retinal imaging biomarkers in AD, amyloid and vascular dysfunction, conveys novel information about 1) the localization of retinal amyloid deposits, which, like AD brain patterns, were mostly detected in the peri-arteriolar as compared to the peri-venular regions, and 2) the relationship between the differentiated perivascular AP counts with the cognitive performance and neuroimaging markers. We found that the secondary vascular branches have significantly higher perivascular amyloid burden in subjects with impaired compared to normal cognition, whether stratified by the neuropsychometric diagnosis, MOCA or CDR, and this remained consistent across sex. Patients with even slightly lower MOCA and greater CDR scores had statistical significantly greater peri-venular amyloid burden, that also exhibited significant and negative correlation with HV. Secondary branch peri-arteriolar and peri-venular AP counts significantly and positively correlated with the WMHI count. These promising exploratory findings encourage future studies in larger cohorts to assess the efficacy of monitoring AD progression, response to therapy, and risk of amyloid-related imaging abnormalities (ARIA), via noninvasive retinal perivascular amyloid imaging.

Brain vascular dysfunction in AD has been studied to evaluate the AD pathophysiology and identify specific therapeutic targets (55–57). Vascular alteration in form of pericyte loss in the brain (3) and retina (16) is linked to a rapid neurodegeneration cascade and amyloid deposition, leading to increased retinal and brain vascular amyloidosis. Blood-retina barrier abnormalities and vascular-associated retinal Aβ deposition were reported in postmortem retinas of AD patients inside the blood vessel walls, around and along blood vessels (11, 42, 48, 49). Furthermore, various blood-retinal barrier tight junction biomarkers were found to be deficient in MCI and AD retina and associated with retinal and brain amyloid burden and cognitive status (49). More Aβ_40_ was found to be accumulated in arterioles than in venules (49). Prior retinal fundus photography studies reported both arteriolar and venular dysfunction (19, 26–29) and OCTA studies showed decreased densities of superficial and deep retinal vascular plexuses in AD (58). Other than the specific pericyte injury, the location of the retinal vascular dysfunction that leads to retinal perivascular amyloid deposition, and if it differentially affects arterioles versus venules at various stages of AD progression, remains currently unclear (59–62).

Similarly, the link between the blood-brain barrier and cerebral neurovascular unit breakdown (63–65) in AD was explored in various studies (7, 66). Increased cerebral capillary damage had been shown to correlate with insoluble Aβ level in AD (67), and pericyte downregulation in the deep white matter was shown in both vascular dementia and AD (68). Aβ deposition around cerebral blood vessels and mostly arteries (CAA), is thought to be major contributor to vascular dysfunction in AD (69–72). The two-hit vascular hypothesis of AD emphasizes the role of neurovascular dysfunction as an early factor that favors vascular Aβ aggregation and neurodegeneration (72): vascular dysfunction (hit one) is followed by the Aβ accumulation (hit two) that promotes and precedes neurodegeneration (73). According to the vascular hypothesis of AD, an alteration in the neurovascular unit could lead to Aβ vascular accumulation that promotes neuronal dysfunction, accelerating neurodegeneration and dementia, and insoluble vascular amyloid deposits trigger neurovascular unit dysfunction in AD brain (74). Whereas most human studies focused on the amyloid deposition on leptomeningeal and cortical arterioles, murine models of AD have also shown abnormal venules (75). Additionally, the brain glymphatic system is impaired in AD, leading to insufficient amyloid clearance (76), and the development of novel perivascular clearance system imaging techniques of the brain are underway (77).

As the phenomenon of vascular amyloid clearance had not been inspected in the retina, we conducted this topographic analysis of retinal fluorescent images to assess the location of AP in relationship to the primary, secondary and tertiary retinal arteriolar and venular branches, in subjects with normal and impaired cognition. Multiple prior studies have shown that retinal proximal mid-periphery has greater accumulation of amyloid compared to the posterior pole (17, 39, 42, 44, 45). In the retinal supero-temporal quadrant, we noted more perivascular amyloid plaque in the peri-arteriolar region when compared to the peri-venular region, overall, as well as in the primary, secondary and tertiary branches. Our discovery that amyloid accumulation in the retina occurs more frequently along the arterioles is consistent with the pattern of cerebral arterial amyloidosis (16, 71, 72). The analysis of the retinal supero-temporal quadrant with relatively high amyloid burden (42, 51) revealed a significant increase in AP accumulation in the peri-venular and peri-arteriolar secondary and tertiary branch regions of subjects with CDR scores of 1 and 2, and MOCA scores lower than 26. Additionally, the perivascular amyloid burden around the secondary small and tertiary branches inversely correlated with HV. The secondary branch peri-arteriolar and peri-venular plaque burden also positively correlated with the number of WMHI lesions.

These findings are hypothesis generating. Venular narrowing and increased venular tortuosity were described in AD, in correlation with the cognitive status (15, 17, 26, 29, 78). It is possible that the overall retinal peri-venular amyloid burden may become a marker of cognitive performance and hippocampal atrophy, whereas secondary branches perivascular AP burden will correlate not only with hippocampal atrophy, but also with WMHI burden. These suggest the possibility that whereas the retinal arteriolar dysfunction may be responsible for the peri-arteriolar amyloid deposition early on, the venular dysfunction may appear later in the disease process and can serve as a discriminator of more advanced disease and cognitive impairment. The association between WMHI and CAA is gaining increased recognition (79). Deposition of Aβ_40_ in the walls of cerebral arteries suggests an age-related failure of perivascular drainage of soluble Aβ from the brain. White matter pathology is suggested to be the link between blood-brain barrier leakage and decline of information processing speed in older individuals with and without cognitive impairment (80). The negative association between white matter blood-brain barrier leakage and information processing speed performance was mediated by the WMHI volume, and leakage did not correlate with the HV. (80). The brain microcirculation disruption not only contributes to amyloidopathy but also initiates a non-amyloidogenic pathway of vascular-mediated neuronal dysfunction and injury, with increased permeability of blood vessels, leakage of blood-borne components into the brain, and, consequently, neurotoxicity. Vascular dysfunction also includes a diminished brain capillary flow, causing multiple focal ischemic or hypoxic microinjuries, diminished Aβ clearance, and formation of neurotoxic oligomers, which lead to neuronal dysfunction (81).

There seems to be greater retinal amyloid accumulation around first-order branches in cognitively intact individuals, and significantly greater retinal amyloid accumulation around smaller branches in patients with impaired cognition. The reduced complexity of the retinal vascular branching network, that was described before in subjects with MCI and AD (15, 26, 29, 82), may explain the preferential secondary branch perivascular amyloid accumulation in cognitively impaired. Interestingly, an agnostic machine learning-based study highlighted the same secondary branch perivascular area on the heat-maps derived from retinal photographs that discriminated between AD and healthy controls with high accuracy (83).

Our preliminary study results will require confirmation in larger studies, which should further explore the clinical usefulness of the perivascular amyloid plaque topography, and the underlying pathophysiology of the peri-arteriolar and peri-venular amyloid deposition in various stages of AD progression. With this current methodology, we could not assess the retinal far periphery nor determine the status of the capillary-adjacent amyloid burden, which deserves future investigation. Our limited number of subjects with cognitive decline, of mainly Caucasian race, and the lack of PET confirmation of their cerebral amyloid status constitute important study limitations that reduce our results’ generalizability.

## Conclusion

In this exploratory study, we found that the accumulation of the amyloid in the distal peri-venular regions of the retina can be used to distinguish between normal and impaired cognition and is inversely related to hippocampal volume. Our results further support the hypothesis that the pathophysiology of AD in the retina and brain is similar, as evidenced by the preferential accumulation of amyloid in the peri-arterial regions. However, the relationship between retinal perivascular amyloid deposition and cognitive performance is complex and requires further investigation in larger longitudinal prospective studies. It is worth exploring the potential of peri-arteriolar and peri-venular amyloid burden as biomarkers of various stages of AD.

## Dedication

The authors dedicate this manuscript to the memory of Dr. Salomon Moni Hamaoui and Lillian Jones Black, both of whom died from Alzheimer’s disease.

## Consent statement

Informed consent was obtained from all subjects involved in the study.

## Data availability statement

Data available upon request due to restrictions, e.g. privacy or ethical.

## Author contributions

OMD, JD, D-TF, YK, MSM, JS, MK-H: designed and performed experiments, collected and analyzed data, created figures, drafted, and edited the manuscript. SRV, KOJ: retinal image storage and processing. OMD, DSS, PDL: cognitive and behavioral testing and imaging data. OMD, JD, D-TF, JAS, PDL, KLB, ROC, MK-H: data interpretation and discussion. OMD, MK-H: responsible for study conception, design, interpretation of data, supervision, and manuscript writing and editing. All authors have read and approved of this manuscript.

## Diversity, Equity, and Inclusion Statement

We are committed to embrace diversity, promote equity, and foster inclusion in this study, to the best of our ability. We included equal number of females and males, and balanced our case and study groups for age, sex, ethnicity, and years of education. Overall, we aimed to advance Alzheimer’s research that is representative, impactful, and beneficial for all individuals affected by this challenging condition.

