## Supplementary materials for "Distinctive retinal peri-arteriolar versus peri-venular amyloid plaque distribution correlates with the cognitive performance"

*equal first co-authors

Supplementary Tables 1 to 3
Supplementary Figures 1 to 4

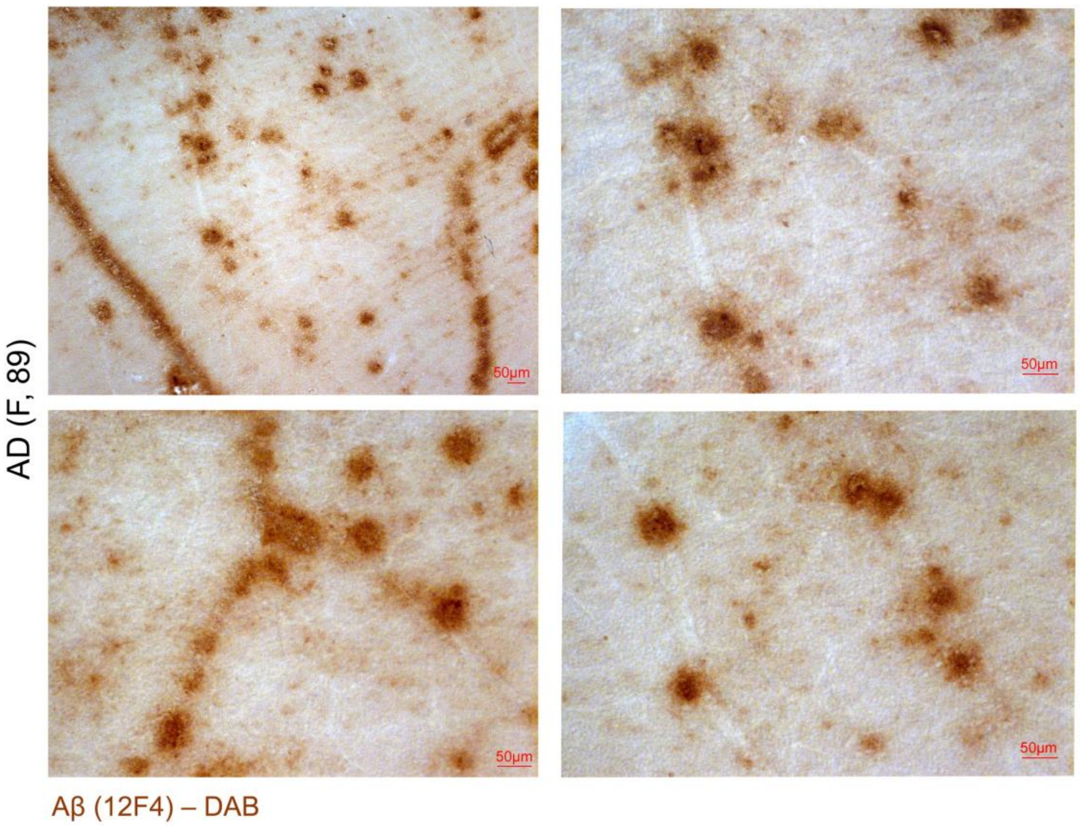


**Supplementary Figure 1. Extended data of histological evidence of retinal vascular and perivascular amyloid deposition.**

The raw representative microscopic images, with no arrow annotations, from a confirmed AD patient shown in Figure 1D.

**Supplementary Table 1. Retinal perivascular amyloid plaques (AP): venules versus arterioles**

| | **Venular AP** | | **Arterial AP** | | **Subject Number** | **Fold Change** | **Paired t test** |
|---|---|---|---|---|---|---|---|
| | Mean | SD | Mean | SD | n | FC | P value |
| **Total Perivascular AP** | **35.57** | 12.66 | **51.96** | 19.65 | 28 | 1.46 | **<0.0001** |
| **Primary Br Perivascular AP** | **13.93** | 10.87 | **16.57** | 17.50 | 28 | 1.19 | 0.34 |
| *Primary Br – main* | ***5.68*** | *5.11* | ***3.25*** | *3.30* | *28* | *0.57* | **0.013** |
| *Primary Br – small* | ***8.27*** | *7.31* | ***14.35*** | *15.21* | *26* | *1.74* | **0.022** |
| **Secondary Br Perivascular AP** | **11.52** | 5.73 | **19.81** | 11.77 | 27 | 1.72 | **0.0013** |
| *Secondary Br – main* | ***4.67*** | *2.87* | ***8.59*** | *4.47* | *27* | *1.84* | **<0.0001** |
| *Secondary Br – small* | ***6.85*** | *4.73* | ***11.22*** | *9.61* | *27* | *1.64* | **0.039** |
| **Tertiary Br Perivascular AP** | **11.58** | 11.58 | **15.13** | 10.23 | 24 | 1.31 | 0.17 |

AP –Amyloid plaques; Br – Branch; SD – Standard deviation. Statistical analysis was established using Student’s t test. *P < 0.05, **P < 0.01, ***P < 0.001, ****P < 0.0001.

**Supplementary Table 2. Retinal perivascular amyloid plaques in males and females**

| SEX | | Males | | | Females | | | Fold Change | Unpaired t test |
|---|---|---|---|---|---|---|---|---|---|
| | Vascular type | Mean | SD | n | Mean | SD | n | FC | P value |
| Perivascular AP | Total | **88.00** | 32.43 | 14 | **87.07** | 25.78 | 14 | 0.99 | **0.93** |
| | Primary Br (total) | **29.29** | 26.24 | 14 | **31.21** | 25.78 | 14 | 1.05 | **0.88** |
| | *Primary Br ─ main* | **8.43** | 6.43 | 14 | **9.43** | 7.93 | 14 | 1.12 | **0.72** |
| | *Primary Br ─ small* | **21.36** | 22.35 | 14 | **21.79** | 17.89 | 14 | 1.02 | **0.96** |
| | Secondary Br (total) | **34.64** | 14.01 | 14 | **26.64** | 14.06 | 14 | 0.77 | **0.14** |
| | *Secondary Br ─ main* | **13.71** | 5.70 | 14 | **12.29** | 7.14 | 14 | 0.90 | **0.56** |
| | *Secondary Br ─ small* | **20.93** | 11.72 | 14 | **14.36** | 9.56 | 14 | 0.69 | **0.12** |
| | Tertiary | **22.29** | 14.18 | 14 | **28.14** | 20.70 | 14 | 1.26 | **0.39** |
| Peri-venular AP | Total | **36.36** | 11.05 | 14 | **34.79** | 14.47 | 14 | 0.96 | **0.75** |
| | Primary Br (total) | **12.79** | 10.76 | 14 | **15.07** | 11.26 | 14 | 1.18 | **0.59** |
| | *Primary Br ─ main* | **5.21** | 4.63 | 14 | **6.14** | 5.68 | 14 | 1.18 | **0.64** |
| | *Primary Br ─ small* | **7.57** | 7.79 | 14 | **8.93** | 6.82 | 14 | 1.18 | **0.63** |
| | Secondary Br (total) | **12.50** | 5.46 | 14 | **9.71** | 6.44 | 14 | 0.78 | **0.23** |
| | *Secondary ─ main* | **4.93** | 2.95 | 14 | **4.38** | 2.87 | 13 | 0.89 | **0.63** |
| | *Secondary Br ─ small* | **7.57** | 5.03 | 14 | **6.08** | 4.44 | 13 | 0.80 | **0.42** |
| | Tertiary | **11.77** | 7.70 | 13 | **11.36** | 15.39 | 11 | 0.97 | **0.93** |
| Peri-arteriolar AP | Total | **51.64** | 23.50 | 14 | **52.29** | 15.78 | 14 | 1.01 | **0.93** |
| | Primary Br (total) | **17.00** | 18.83 | 14 | **16.14** | 16.76 | 14 | 0.95 | **0.90** |
| | *Primary Br ─ main* | **3.21** | 3.09 | 14 | **3.29** | 3.60 | 14 | 1.02 | **0.96** |
| | *Primary Br ─ small* | **16.08** | 17.07 | 12 | **12.86** | 13.88 | 14 | 0.80 | **0.60** |
| | Secondary Br (total) | **22.14** | 13.28 | 14 | **16.93** | 9.53 | 14 | 0.76 | **0.24** |
| | *Secondary Br ─ main* | **8.79** | 3.91 | 14 | **8.21** | 5.01 | 14 | 0.93 | **0.74** |
| | *Secondary Br ─ small* | **13.36** | 11.29 | 14 | **8.71** | 6.89 | 14 | 0.65 | **0.20** |
| | Tertiary | **12.23** | 9.72 | 13 | **19.21** | 10.30 | 14 | 1.57 | **0.08** |

AP – Amyloid plaque; Br – Branch; SD – Standard deviation. Statistical analysis was established using Student's t test.

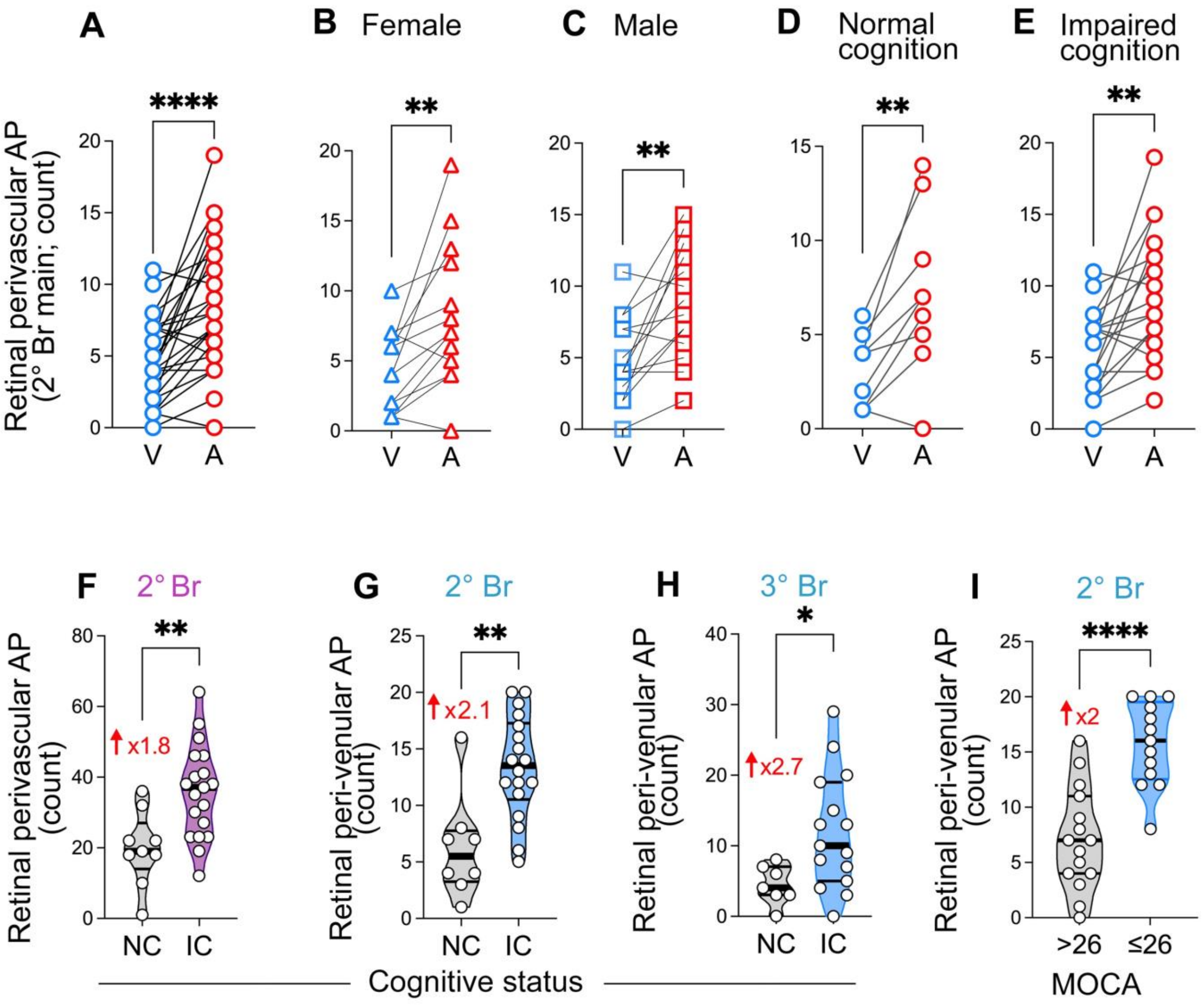


**Supplementary Figure 2. Extended data on retinal perivascular amyloid plaque distribution and stratification by cognitive status and MOCA score.**

(**A-E**) Quantitative analyses of retinal AP count stratified by venules (V) versus arterioles (A) in the total perivascular secondary main branches (**A**), in females (**B**), males (**C**), normal cognition (**D**) and impaired cognition (**E**) groups. (**F-I**) Quantitative analyses of retinal AP counts stratified by cognitive status, NC versus IC (**F-H**) and MOCA score (**I**). Violin plots are showing individual data points, median and interquartile range. * $p<0.05$, ** $p<0.01$, **** $p< 0.0001$ by paired and unpaired two-tailed Student's t test. AP, Amyloid plaque; Br, branches; IC, Impaired cognition; NC, Normal cognition; V, Venule; A, Arteriole; 2° Br, secondary branch; 3° Br, tertiary branch.

**Supplementary Table 3. Retinal perivascular amyloid plaques in patients defined by CDR score.**

| **CDR SCORE** | | **CDR = 0.5** | | | **CDR = 1** | | | **CDR = 2** | | | **Fold Change** | | | **ANOVA** | **Tukey's Multiple Comparison** | | |
|---|---|---|---|---|---|---|---|---|---|---|---|---|---|---|---|---|---|
| | **Vascular type** | Mean | SD | n | Mean | SD | n | Mean | SD | n | 0.5 vs 1 FC | 0.5 vs 2 FC | 1 vs 2 FC | P value | 0.5 vs 1 | 0.5 vs 2 | 1 vs 2 |
| **Perivascular AP** | **Total** | **86.09** | 22.38 | 11 | **76.77** | 22.56 | 13 | **134.00** | 43.52 | 3 | 0.89 | 1.38 | 1.55 | **0.040** | 0.62 | 0.12 | **0.032** |
| | **Primary Br (total)** | **30.64** | 21.63 | 11 | **33.86** | 29.29 | 14 | **33.25** | 39.84 | 3 | 1.11 | 0.47 | 0.42 | 0.50 | 0.95 | 0.60 | 0.46 |
| | *Primary Br – main* | **9.55** | 6.49 | 11 | **8.64** | 7.75 | 14 | **13.00** | 12.27 | 3 | 0.91 | 0.84 | 0.93 | 0.93 | 0.95 | 0.94 | 0.99 |
| | *Primary Br – small* | **21.09** | 16.53 | 11 | **25.21** | 23.12 | 14 | **20.25** | 28.38 | 3 | 1.20 | 0.30 | 0.25 | 0.34 | 0.86 | 0.50 | 0.31 |
| | **Secondary Br (total)** | **28.64** | 14.36 | 11 | **27.71** | 11.53 | 14 | **58.50** | 16.82 | 3 | 0.97 | 1.80 | 1.86 | **0.020** | 0.98 | **0.027** | **0.018** |
| | *Secondary Br – main* | **12.55** | 5.65 | 11 | **11.93** | 6.67 | 14 | **23.25** | 8.265 | 3 | 0.95 | 1.57 | 1.65 | 0.16 | 0.97 | 0.20 | 0.14 |
| | *Secondary Br – small* | **16.09** | 11.38 | 11 | **15.79** | 8.91 | 14 | **35.25** | 11.27 | 3 | 0.98 | 1.99 | 2.03 | 0.051 | 1.00 | 0.060 | **0.048** |
| | **Tertiary** | **25.91** | 14.87 | 11 | **18.79** | 11.76 | 14 | **42.25** | 30.94 | 3 | 0.73 | 2.03 | 2.80 | **0.0058** | 0.47 | **0.028** | **0.0041** |
| **Peri-venular AP** | **Total** | **33.36** | 10.37 | 11 | **32.57** | 9.87 | 14 | **63.50** | 15.80 | 3 | 0.98 | 1.73 | 1.77 | **0.0025** | 0.98 | **0.004** | **0.002** |
| | **Primary Br (total)** | **14.45** | 7.93 | 11 | **14.36** | 12.79 | 14 | **10.25** | 11.35 | 3 | 0.99 | 0.69 | 0.70 | 0.81 | 1.00 | 0.82 | 0.82 |
| | *Primary Br – main* | **6.45** | 4.06 | 11 | **5.21** | 5.89 | 14 | **5.50** | 5.20 | 3 | 0.81 | 0.77 | 0.96 | 0.82 | 0.83 | 0.91 | 1.00 |
| | *Primary Br – small* | **8.00** | 5.53 | 11 | **9.14** | 8.48 | 14 | **4.75** | 6.40 | 3 | 1.14 | 0.63 | 0.55 | 0.68 | 0.92 | 0.81 | 0.66 |
| | **Secondary Br (total)** | **10.11** | 6.41 | 9 | **11.00** | 5.80 | 13 | **26.75** | 22.97 | 3 | 1.09 | 1.52 | 1.39 | 0.39 | 0.83 | 0.35 | 0.54 |
| | *Secondary Br – main* | **3.89** | 2.47 | 9 | **4.15** | 2.38 | 13 | **9.25** | 8.22 | 3 | 1.07 | 1.37 | 1.28 | 0.68 | 0.97 | 0.66 | 0.74 |
| | *Secondary Br – small* | **6.30** | 4.47 | 10 | **6.57** | 5.32 | 14 | **17.50** | 15.00 | 3 | 1.04 | 1.59 | 1.52 | 0.49 | 1.0 | 0.5 | 0.5 |
| | **Tertiary** | **10.56** | 6.15 | 9 | **7.17** | 6.78 | 12 | **26.50** | 19.55 | 3 | 0.68 | 3.06 | 4.51 | **0.0007** | 0.65 | **0.0028** | **0.0005** |
| **Peri-arteriolar AP** | **Total** | **52.73** | 16.19 | 11 | **49.43** | 21.59 | 14 | **70.50** | 28.77 | 3 | 0.94 | 1.16 | 1.23 | 0.66 | 0.91 | 0.80 | 0.64 |
| | **Primary Br (total)** | **16.18** | 16.58 | 11 | **19.50** | 19.49 | 14 | **23.00** | 37.37 | 3 | 1.21 | 0.27 | 0.22 | 0.41 | 0.89 | 0.56 | 0.38 |
| | *Primary Br – main* | **3.09** | 3.83 | 11 | **3.43** | 3.28 | 14 | **7.50** | 9.026 | 3 | 1.11 | 0.97 | 0.88 | 0.96 | 0.97 | 1.00 | 0.98 |
| | *Primary Br – small* | **14.4** | 12.96 | 10 | **16.07** | 17.36 | 14 | **20.67** | 32.35 | 3 | 1.12 | 0.14 | 0.12 | 0.49 | 0.96 | 0.56 | 0.46 |
| | **Secondary Br (total)** | **15.70** | 5.85 | 10 | **16.50** | 9.36 | 14 | **31.75** | 11.79 | 3 | 1.05 | 2.31 | 2.20 | **0.0021** | 0.97 | **0.0023** | **0.0024** |
| | *Secondary Br –main* | **8.45** | 2.88 | 11 | **7.29** | 4.70 | 14 | **14** | 3.46 | 3 | 0.86 | 1.70 | 1.97 | **0.036** | 0.75 | 0.08 | **0.028** |
| | *Secondary Br – small* | **7.60** | 5.32 | 10 | **9.21** | 7.15 | 14 | **17.75** | 12.53 | 3 | 1.21 | 2.89 | 2.39 | **0.014** | 0.84 | **0.01** | **0.02** |
| | **Tertiary** | **17.27** | 10.47 | 11 | **13.62** | 9.30 | 13 | **15.75** | 16.48 | 3 | 0.79 | 1.18 | 1.49 | 0.53 | 0.68 | 0.90 | 0.59 |

AP – Amyloid plaque; Br – Branch; CDR – Clinical dementia rating; SD – Standard deviation. Statistical analysis was established using one-way ANOVA and Tukey's multiple comparison post-test. *P < 0.05, **P < 0.01, ***P < 0.001, ****P < 0.0001

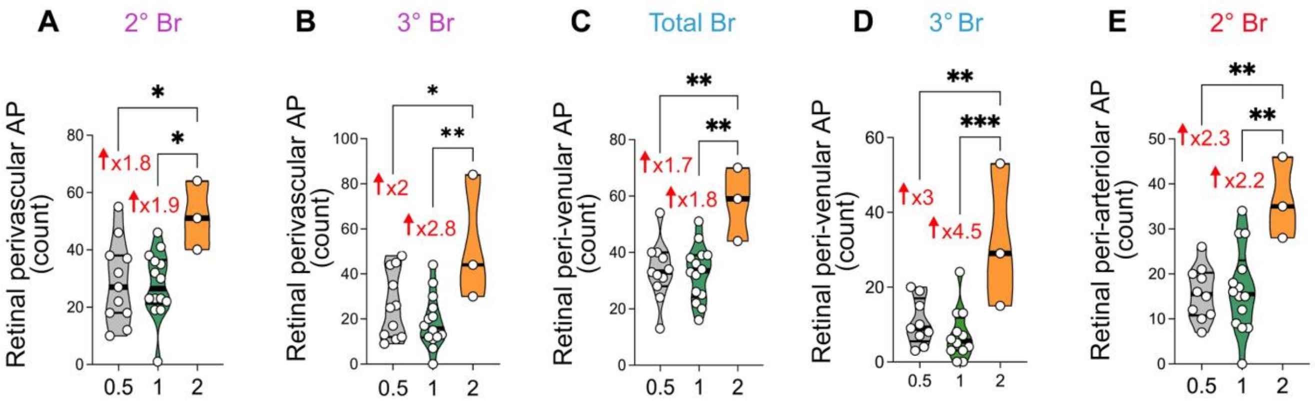


**Supplementary Figure 3. Retinal perivascular amyloid plaque count stratified by CDR.**
(**A-E**) Quantitative analyses of retinal AP count in total perivascular secondary (**A**) and tertiary branches (**B**), in total peri-venular branches (**C**) and tertiary branches (**D**), and total peri-arteriolar secondary branches. Violin plots are showing individual data points, median and interquartile range. * $p<0.05$, ** $p<0.01$, *** $p< 0.001$ by one-way ANOVA followed by Tukey's multiple comparisons test. AP, Amyloid plaques; 2° Br, secondary branch; 3° Br, tertiary branch.

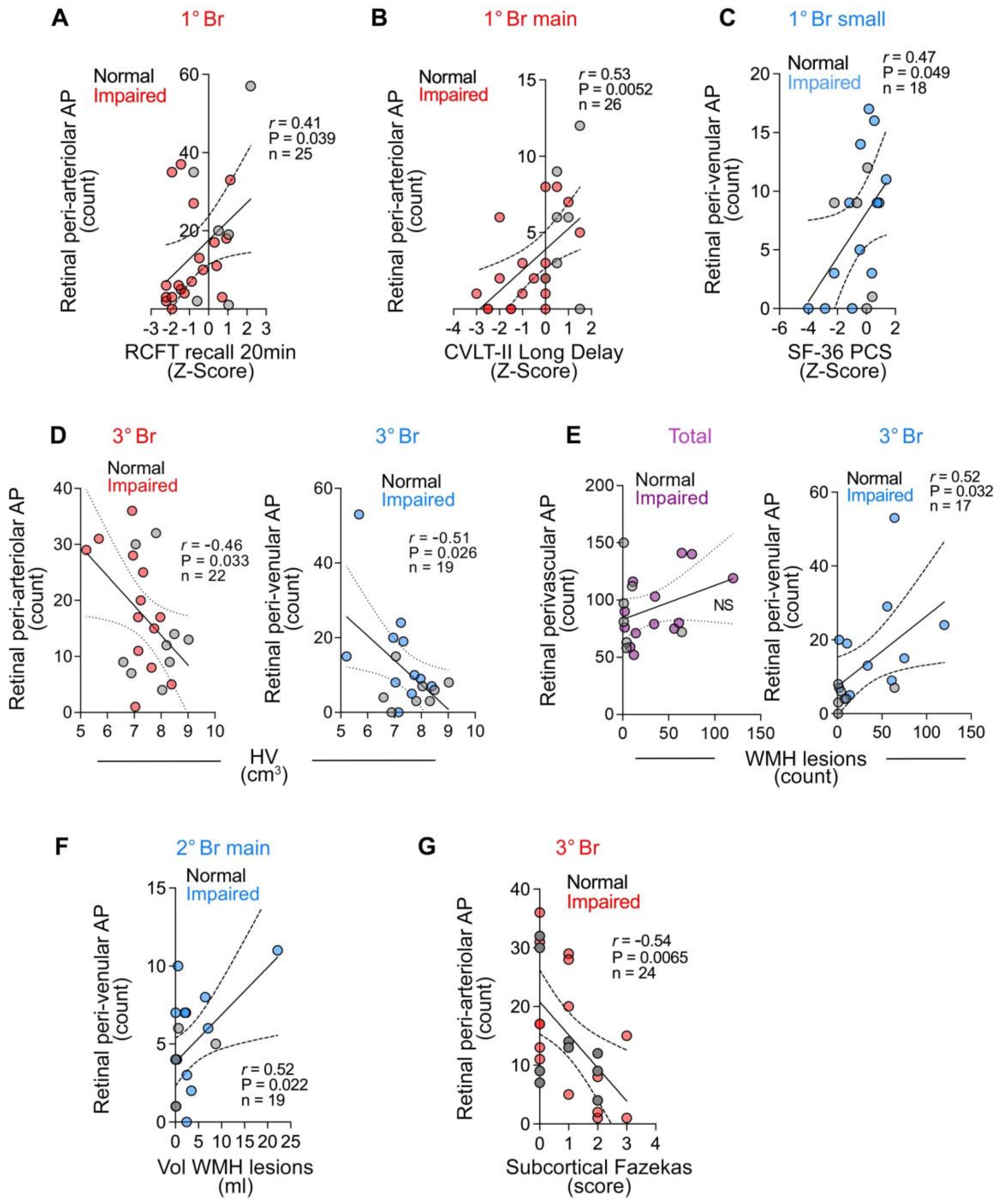


**Supplementary Figure 4. Additional correlation analyses between retinal perivascular AP distribution and cognitive and neuroimaging measures.**

Pearson's *r* correlation analyses between retinal AP count and RCFT-recall 20min (**A**), CVLT-II Long delay (**B**), SF-36 PCS (**C**), hippocampal volume (**D**), number (**E**) and volume (**F**) of white matter hyperintensities lesions, and Subcortical Fazekas (**G**). AP, Amyloid plaques; 1° Br, primary branch; 2° Br, secondary branch; 3° Br, tertiary branch.
